# Integrated molecular analysis reveals two distinct subtypes of pure seminoma of the testis

**DOI:** 10.1101/2022.04.25.489437

**Authors:** Kirill E. Medvedev, Anna V. Savelyeva, Aditya Bagrodia, Liwei Jia, Nick V. Grishin

**Affiliations:** Department of Biophysics, University of Texas Southwestern Medical Center, Dallas, TX 75390, USA; Department of Urology, University of Texas Southwestern Medical Center, Dallas, TX 75390, USA; Department of Urology, University of California San Diego Health, La Jolla, CA 92037, USA; Department of Pathology, University of Texas Southwestern Medical Center, Dallas, TX 75390, USA; Department of Biochemistry, University of Texas Southwestern Medical Center, Dallas, TX 75390, USA

## Abstract

Testicular germ cell tumors (TGCT) are the most common solid malignancy in adolescent and young men, with a rising incidence over the past 20 years. Overall, TGCTs are second in terms of the average life years lost per person dying of cancer, and clinical therapeutics without adverse long-term side effects are lacking. Platinum-based regimens for TGCTs have heterogeneous outcomes even within the same histotype that frequently leads to under- and over-treatment. Understanding of molecular differences that lead to diverse outcomes of TGCT patients may improve current treatment approaches. Seminoma is the most common subtype of TGCTs, which can either be pure or present in combination with other histotypes. Here we conducted a computational study of 64 pure seminoma samples from The Cancer Genome Atlas, applied consensus clustering approach to their transcriptomic data and revealed two clinically relevant seminoma subtypes: seminoma subtype 1 and 2. Our analysis identified significant differences in pluripotency stage, activity of double stranded DNA breaks repair mechanisms, rates of loss of heterozygosity, DNA methylation, and expression of lncRNA responsible for cisplatin resistance between the subtypes. Seminoma subtype 1 is characterized by higher pluripotency state, while subtype 2 showed attributes of reprogramming into non-seminomatous TGCT. The seminoma subtypes we identified may provide a molecular underpinning for variable responses to chemotherapy and radiation. Translating these findings into clinical care may help improve risk stratification of seminoma, decrease overtreatment rates, and increase long-term quality of life for TGCT survivors.

## Introduction

Testicular germ cell tumor (TGCT) is the most common solid cancer among men aged 15-44 years [1]. Two major types of TGCTs are seminomatous (SE) and non-seminomatous (NSE) [2]. NSE TGCTs include embryonal carcinoma (EC), teratoma (TE), yolk sac tumor (YST), choriocarcinoma and mixed GCTs, which include combinations of any NSE or SE subhistologies. Mixed type of TGCTs in its own turn represents the most common type of NSE, since pure EC, TE and YST are rare [3]. However, seminoma is the most common histological subtype of TGCT among young men 15-44 years of age [1,4,5]. In 2021 the American Cancer Society estimated around 9,470 TGCT cases in the US [6] and around 39,000 cases worldwide [7]. TGCT is the second cancer type (first being all pediatric cancers combined) based on calculation of life years lost per person dying of cancer [8]. NSE tumors are more aggressive compared to SE, requiring more intensive treatment approaches [8–11].

Management of patients with seminoma starts with orchiectomy followed by observation, platinum-based chemotherapy (cisplatin) or radiation therapy [12–14]. Despite a high patient survival rate, current treatments significantly decrease patients’ quality of life and can cause combinations of around 40 severe adverse long-term side effects like infertility, neurotoxicity, hypercholesterolaemia, secondary cancers, and death [15–17]. Following chemotherapy, TGCT patients demonstrated a 3.6-dB decline in hearing for every 100mg/m^2^ increase in cumulative cisplatin dose [18–19]. BEP-treated patients received more than 400mg/m^2^ of cisplatin had impaired renal function at the end of treatment and 20% decrease of glomerular filtration rate after 5 years of follow-up [20–21]. Moreover, about 20% of seminoma patients will experience relapse, and the reason for this phenomenon is unclear [22–23]. Relapsed patients will be treated again using conventional and high-dose chemotherapy with stem cell transplant that aggravate side effects drastically.

Multiple clinical studies demonstrate heterogeneous patient outcomes in the treatment of seminoma patients. [24–26] Despite limited understanding of seminoma intratumoral heterogeneity [27–28], the existence and relevance of seminoma subtypes remains unclear and has never been studied in detail.

Recent progress in understanding the molecular heterogeneity of cancer types and intertumoral heterogeneity has suggested improvements for therapeutic strategies [29–30]. A large variety of cancer types such as meningioma [31], pancreatic neuroendocrine tumors [32], and squamous cell carcinoma of the head and neck [33] reveal the existence of clinically relevant subtypes with distinctive molecular characteristics. Moreover, clinically relevant subtypes for breast [34–35] and lung cancer [36] have led to different treatment strategies in the clinic, which highlights the importance of subtype-specific therapy. A better understanding of cancer heterogeneity and the identification of subtypes with distinctive clinical characteristics should lay the basis for future applications of personalized cancer therapy aimed to increase the efficiency of patient treatment with reduced toxicity and side effects [37]. Various methods for precision cancer medicine are used in clinics for management of cancer patients. Amongst them are diagnostic methods based on genome profiling [38], analysis of tumor proteomic [39] and transcriptomic [40] data, which have been successfully applied to the most abundant cancer types including lung [41], breast [42], and prostate [43] cancers.

Here we conducted a computational study of 64 pure seminoma samples from the TCGA data portal. We applied consensus clustering approach to transcriptomic data that revealed two distinct seminoma subtypes. Analysis of transcriptomic, genomic and epigenomic data showed similarity of seminoma subtype 2 with non-seminomatous GCTs. Therefore, we propose that consideration of identified subtypes might help to improve seminoma clinical management by administrating subtype-specific treatment.

## Materials and Methods

### Data collection

The RNA-sequencing data (HTSeq-count), histopathological slides, DNA methylation data, copy number variation data, single-nucleotide variants, level of lymphocytes infiltration and corresponding patient clinical information (race, ethnicity, clinical stage) were collected for 64 pure seminoma cases (TCGA-TCGT project) from The Cancer Genome Atlas (TCGA) data portal (https://portal.gdc.cancer.gov). 162 available histopathological slides were evaluated by a pathologist subspecialized in surgical oncology and genitourinary pathology (L.J.). Two cases (TCGA-2G-AAG9 and TCGA-2G-AAH0) were removed from our dataset due to additional types of TGCT (teratoma and embryonal carcinoma) identified on the slides. Two other cases (TCGA- 2G-AAFG and TCGA-2G-AAHP) include primary and secondary tumors that were considered as different cases with additional number after case ID, ‘1’ for primary and ‘2’ for secondary tumor respectively (for example TCGA-2G-AAFG1 and TCGA-2G-AAFG2). Telomer lengths data for pure seminoma TCGA cases was retrieved from previous study [44]. List of long non-coding RNAs was retrieved from LNCipedia version 5.2 [45].

### Data processing and consensus clustering

Pure seminomas are commonly infiltrated by lymphocytes [46]. To focus on tumor transcriptome, we incorporated an additional filtration step that removes immune cell transcripts (filtered gene set). Immune cell transcripts were defined using the Database of Immune Cell Expression [47]. ConsensusClusterPlus R-package [48] was used to identify transcriptional clusters on the filtered gene set. We used 1000 iterations, 80% sample resampling from 2 to 6 clusters (k2 to k6) using hierarchical clustering with average innerLinkage and finalLinkage, as well as Spearman correlation as the similarity metric. Clustering significance was checked in pairwise comparisons using SigClust [49]. Gene expression data was median centered and log2 transformed. The R function *hclust* was used for unsupervised hierarchical clustering of pure seminoma samples with Ward’s method, and the resulting heatmap was cut with the R function *cutree*. Boxplots were generated using R function *ggplot*. Comparison p-values for boxplots were calculated using Wilcoxon test. For epigenetic data we calculated the standard deviation (SD) of beta values across the all pure seminoma samples for each CpG probe. Autosomal CpG probes with an SD of greater than 0.2 (n=6,208) were used for analysis.

### Identification of differentially expressed genes in seminoma subtypes

Differentially expressed genes for identified seminoma subtypes were obtained using DESeq2 package [50] and raw counts from TCGA as the input data. DESeq2 parameters were set by default. At the first step we selected genes with baseMean > 10 (non-filtered gene set) and genes with baseMean > 5 (filtered gene set). Then, we used Log2 Fold Change > 2 and adjusted P-value < 0.005 to identify genes with significant differential expression. Signature genes differentially expressed in different types of TGCTs were retrieved from previous study [51].

### Gene Set Enrichment Analysis (GSEA)

GSEA [52] was employed to identify characteristically molecular pathways enriched or depleted in the seminoma subtypes. We created a ranked list of genes with distinct level of expression between 2 seminoma subtypes. For that we processed the full list of 53644 transcripts and removed those that do not have gene names. Then, we removed genes for which all samples have zero counts or extreme count outlier or low mean normalized count (automatic DESeq2 filtering). As the output we generated a list of 14477 genes. For each of the genes we calculated a rank using formula: (sign of Log2FC)*(-Log10(adjusted p-value)). Ranked gene list was uploaded to GSEA program. The reference gene set “h.all.v7.4.symbols.gmt [Hallmarks]” was obtained from the Molecular Signatures Database (MSigDB). The number of gene set permutations were 1000 times for each analysis. Groups with minimum 5 and maximum 500 genes were selected. The FDR q-value < 0.05 and normalized enrichment score (NES) > 1.5 were considered significant. Top 1000 DEGs were selected based on the gene rank.

### Loss of heterozygosity

Loss of heterozygosity (LOH) data for all pure seminoma cases was downloaded from TCGA PanCanAtlas study [53]. Difference between subtypes LOH was evaluated using Chi-squared test. P-value < 0.05 was considered significant.

### Biomarkers identification

Potential biomarkers were defined among genes differentially expressed between two identified seminoma subtypes (adjusted P-value < 0.005, |Log2FC| > 1). Biomarkers were rated by the specificity (true negative rate). The specificity was calculated based on the level of genes expression using the minimal value for one of subtypes as a threshold. Pseudogenes and novel transcripts were removed from consideration.

## Results

### Transcriptomic data analysis reveals two distinct molecular subtypes of pure seminomas

Overall, 64 pure seminoma cases from TCGA database were used for this study. Consensus average linkage hierarchical clustering of 64 samples identified two distinct transcriptomic seminoma subtypes (Fig. 1). Consensus matrix heatmap is shown on Fig. S1. The detailed clinical and pathological information for subtypes is summarized in Table 1. There was no notable difference at the patient demographics or tumor stage distribution between the two subtypes.

**Figure 1.**
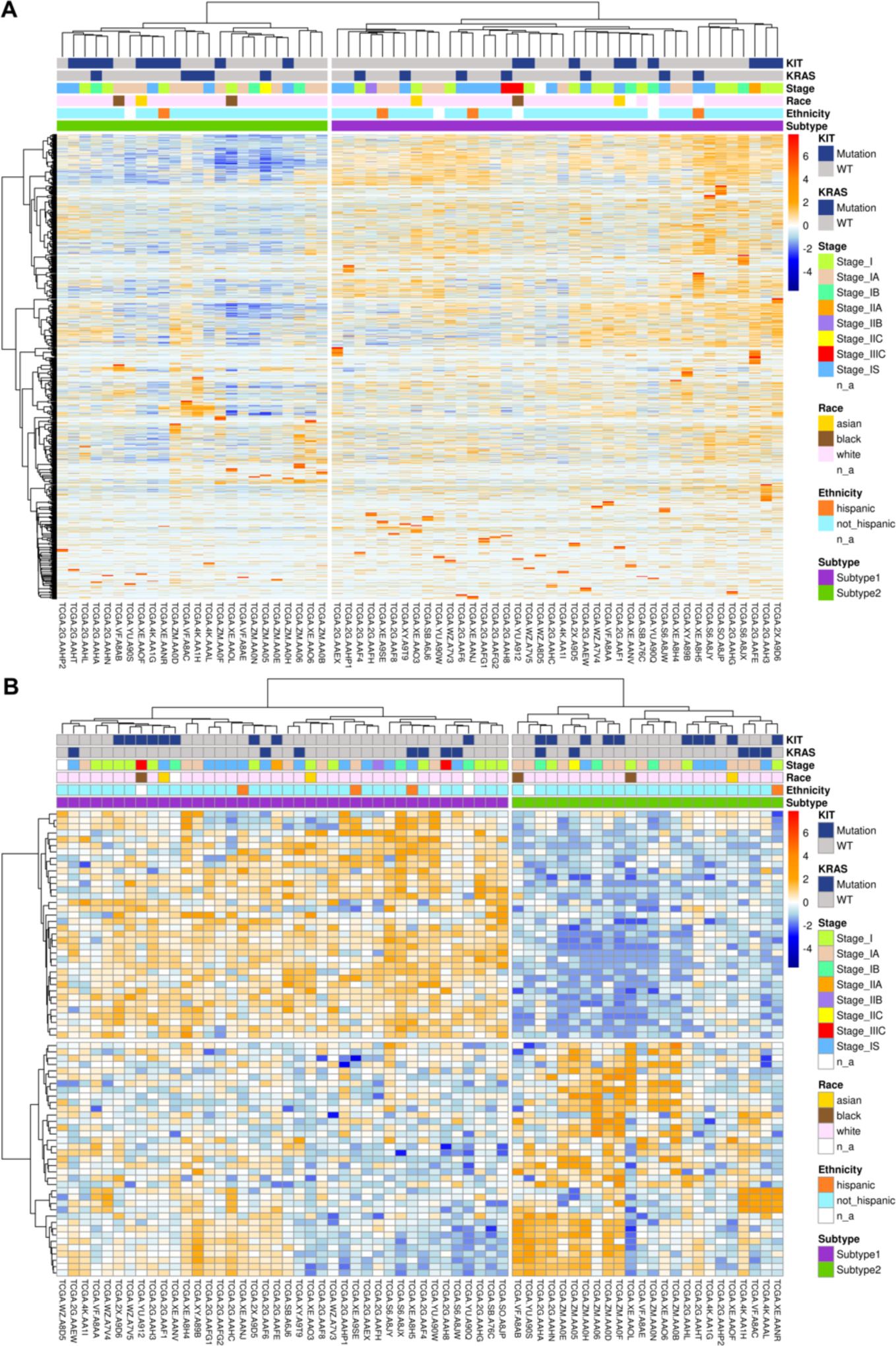
**(A)** Unsupervised hierarchical clustering of seminoma samples based on transcriptomic data. **(B)** Differentially expressed genes between two seminoma subtypes.

**Table 1.** Subtype-based clinical data of seminoma patients.

|  | Subtype 1<br>(total = 40) | Subtype 2<br>(total = 24) |
| --- | --- | --- |
| <u>Average age</u> | 33.03 | 35.75 |
| <u>KIT mutations</u> | 9 | 10 |
| <u>KRAS mutations</u> | 7 | 5 |
| <u>Stage</u> |  |  |
| I | 12 | 4 |
| IA | 7 | 11 |
| IB | 3 | 3 |
| IS | 13 | 5 |
| IIA | 1 | 0 |
| IIB | 1 | 0 |
| IIC | 0 | 1 |
| IIIC | 2 | 0 |
| n/a | 1 | 0 |
| <u>Race</u> |  |  |
| white | 34 | 21 |
| black | 1 | 2 |
| asian | 2 | 1 |
| n/a | 3 | 0 |
| <u>Ethnicity</u> |  |  |
| hispanic | 3 | 1 |
| not hispanic | 34 | 22 |
| n/a | 3 | 1 |

### Two revealed seminoma subtypes differ in pluripotency state and utilized mechanisms of double stranded DNA breaks (DSB) repair

To explore key molecular features of the identified seminoma subtypes, we conducted analysis of differentially expressed genes (DEGs) using the filtered gene set (without B- and T-immune cell transcripts). We found 73 DEGs up- and down-regulated between two seminoma subtypes (Fig. 1B; Supplementary Table S1). Removing immune cell transcripts is rather stringent criterion helping to avoid potential biases of hierarchical clustering related to immune infiltration of seminomas. However, some of the T- and B- immune cell transcripts are also expressed in tumor cells, so we used the non-filtered set of genes for the remaining analysis. We also conducted DEGs analysis on the whole (non-filtered) set of genes, and generated a list of 229 genes with significant differential expression (Supplementary Table S2).

To identify active molecular pathways characterizing either of 2 seminoma subtypes, we analyzed expression of hallmark gene sets [54]. We applied GSEA to a pre-ranked list of DEGs (ranking based on the adjusted p-value and a sign of Log2FC). GSEA revealed 3 gene sets for subtype 1 and 21 gene sets for subtype 2 with normalized enrichment score > 1.50 and FDR q-value < 0.05 (Fig. 2A). All 3 gene sets detected for subtype 1 play an important role in cell cycle progression: mitotic spindle assembly, G2/M cell cycle transition, G1/S phase progression controlled by E2F transcription factors. We added additional steps to GSEA analysis of subtype 2 allowing us to focus on gene sets with the most prominent activation. We calculated a ratio of top 1000 DEGs per each of the revealed gene sets and selected top 7 gene sets with the ratio > 10% (Supplementary table S3, tab ‘GSEA’). Those 7 gene sets fell in 3 major categories: metabolism, immune response and DNA damage response. The metabolism group was represented by genes activated by reactive oxygen species (ROS), genes encoding proteins involved in oxidative phosphorylation (OXPHOS), and genes involved in processing of drugs and other xenobiotics. Immune response gene sets included genes up-regulated during allograft rejection and genes regulated by NF-kB in response to tumor necrosis factor (TNF). DNA damage response group comprised of gene sets associated with DNA reparation and p53 signaling. Next, for selected 10 gene sets (3 for subtype 1 and 7 for subtype 2), we conducted a functional analysis of genes represented in top 1000 DEGs (Fig. 2B). We have found that for subtype 1, genes BRCA2 and SMC1A were related to all 3 detected gene sets. Both, BRCA2 and SMC1A, are key players of homologous recombination (HR) DNA repair. Moreover, more than a half of top 1000 DEGs related to E2F gene set (Fig. 2B, red text) participate in HR repair. We revised a full list of subtype 1 DEGs associated with E2F gene set and found additional well-known players of HR repair (Supplementary table S3, tab “E2F”, green). We compiled a list of detected HR repair genes in subtype 1 (BRCA2, STAG1, SMC1A, SMC3, SMC4, SMC6, MCM2, MCM4, MCM7, RAD21) and populated it with RAD51 and BRCA1 – key mediators of HR repair. The list was used to generate a heatmap of HR repair gene expression in analyzed seminoma samples (Fig. 3A). It is notable that the majority of subtype 1 seminomas were characterized by higher activity of HR repair genes in comparison to subtype 2 seminomas. At the same time subtype 2 samples characterized by increased expression level of genes associated with p53 signaling, and p53 is implicated in multiple repair pathways including DSB repair via c-NHEJ [55, 56].

**Figure 2.**
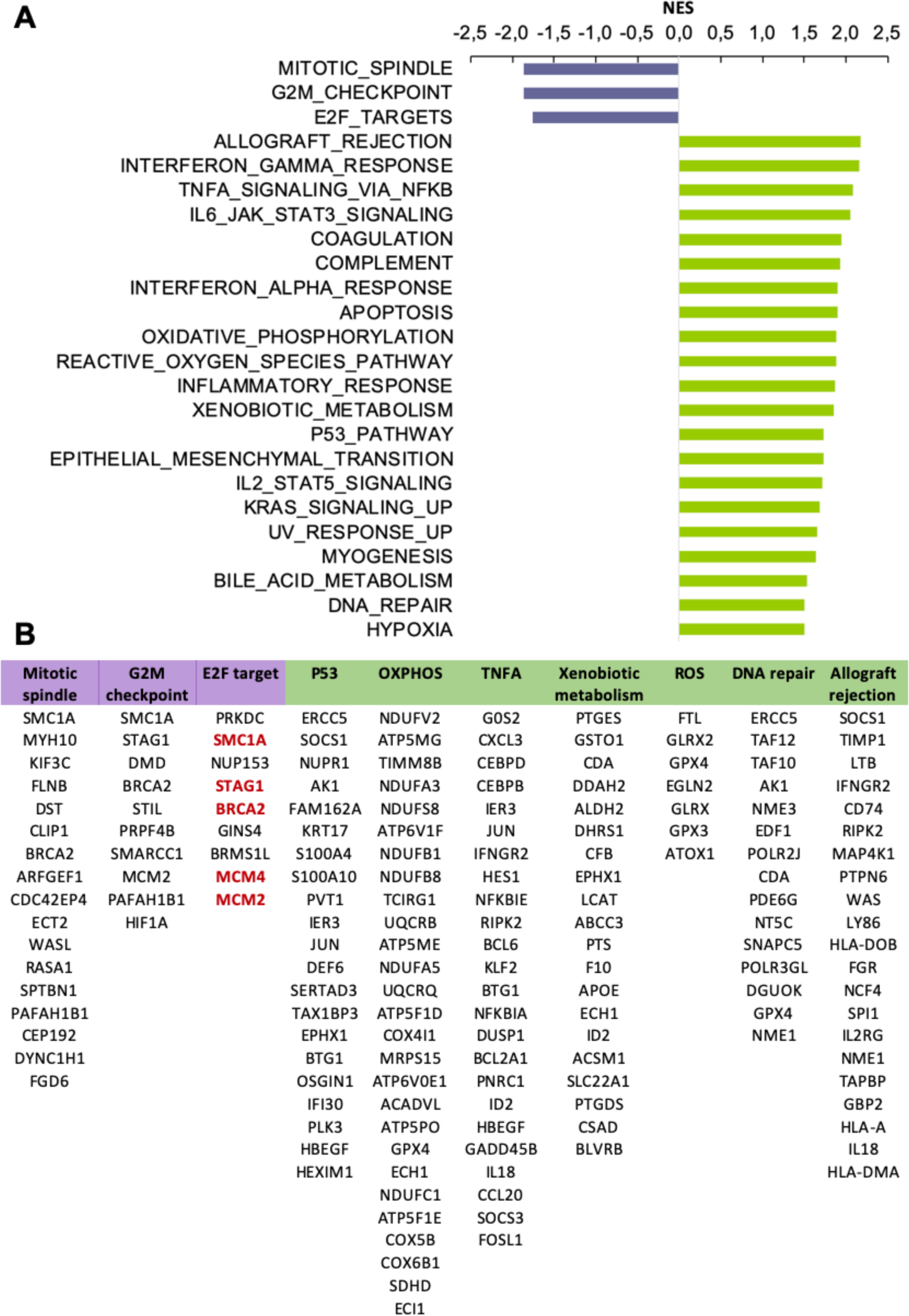
**(A)** Top gene sets enriched in seminoma subtype 1 (purple) and 2 (green) based on the gene set enrichment analysis, NES >1.5, FDR p-value < 0.05. **(B)** Genes from top 1000 DEGs list related to key gene sets enriched in seminoma subtypes. HR-repair genes highlighted in red.

**Figure 3.**
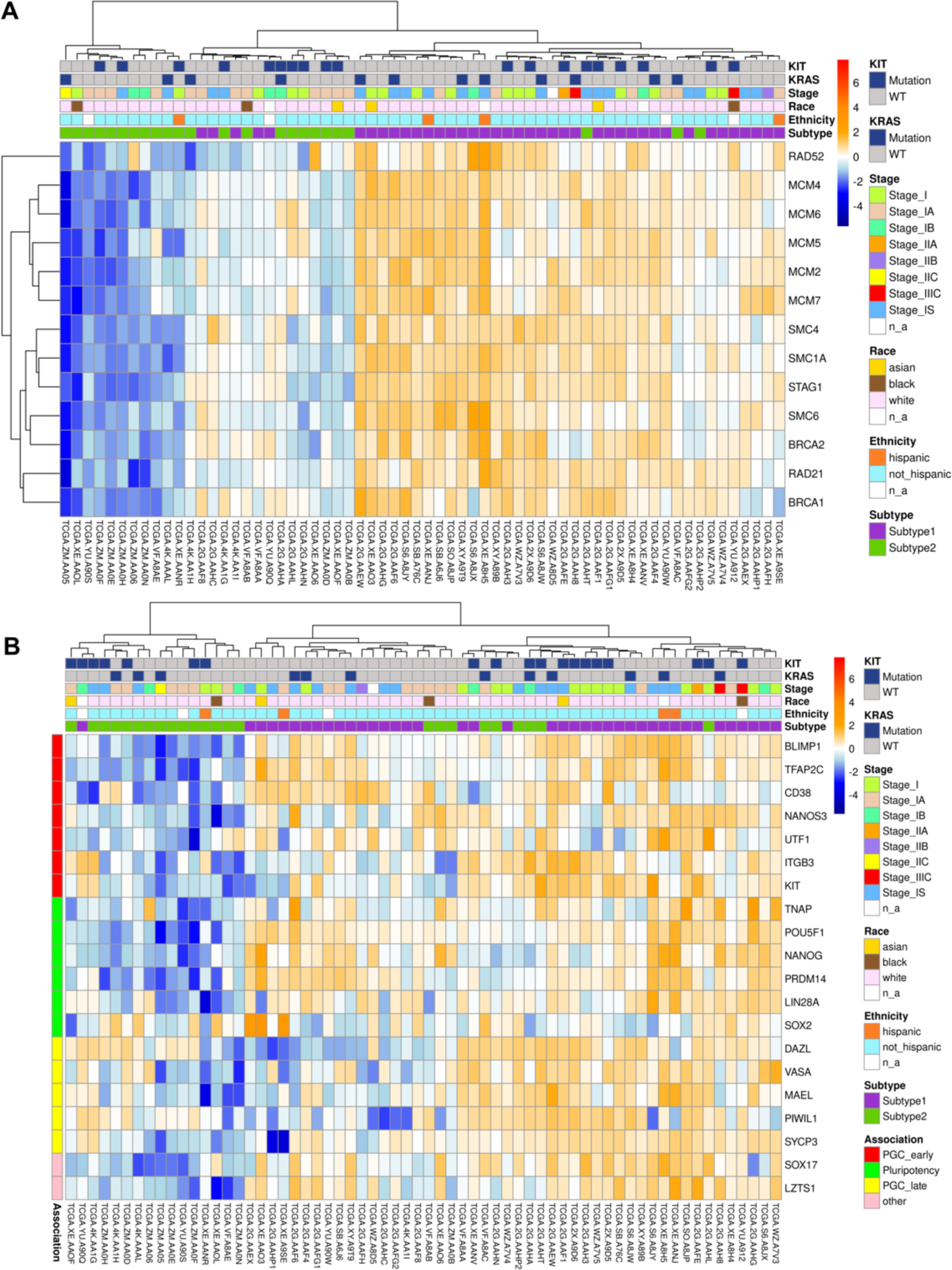
**(A)** Heatmap of expression for HR-mediated DNA repair genes in seminoma subtypes. **(B)** Pluripotency stage of revealed seminoma subtypes. The heatmap shows expression of biomarkers of pluripotency (green), early PGC (red), late PGC (yellow), and mediators of PGC specification (pink, SOX17 and LZTS1).

HR repair as well as classical non-homologous end joining (c-NHEJ) repair are two key mechanisms utilized by cells to defend against double stranded DNA breaks (DSBs) [57]. In general, HR repair is more active during S- and G2-phases as it requires a homologous DNA sequence of the sister chromatid. Classical NHEJ is a faster, but error prone process that is active throughout the cell cycle, and is dominant in G0- or G1-phases. The balance between HR and c-NHEJ repair activation in the response to DSBs depends on the type and location of the cell [58]. Differentiation of inducible pluripotent or embryonic derived stem cells leads to impairment of DNA damage repair via HR repair and has no effect on c-NHEJ [59]. Therefore, high activity of HR repair may be a reflection of a cell stemness status. Group of testicular germ cell cancers unites variety of tumor histological types at the different stages of differentiation, from the most pluripotent embryonal carcinoma to highly differentiated teratoma [60]. Primordial germ cell (PGC) considered as a cell of origin for testicular germ cell tumors (TGCTs) [61]. However, there are several evidences that significant difference in histology and pluripotency state of TGCT subtypes is related to the stem cell hierarchy stage of the initiating cell [62, 63]. We hypothesized that revealed differences in activity of DSB repair mechanisms maybe related to the differentiation state of the seminoma subtypes. To test that, we built a heatmap of expression for key genes associated with early primordial germ cells (PGCs) (BLIMP1, TFAP2C, DND1, CD38, NANOS3, UTF1, ITGB3, KIT), late PGCs (DAZL, VASA, MAEL, PIWIL, SYCP3), as well as pluripotency (TNAP, POU5F1, NANOG, PRDM14, LIN28A, SOX2) [64] (Fig. 3B). The heatmap showed that the majority of subtype 1 tumors had elevated level of pluripotency markers in comparison to subtype 2. This finding supports our theory of seminoma subclassification based on the pluripotency state.

On the contrary, subtype 2 of seminoma lacked both, expression of pluripotency and early PGC markers, and had minimal expression of late PGC markers. This shows that subtype 2 seminomas may be at an advanced differentiation stage. Another piece of evidence supporting this is noticeably higher expression level of ROS- and OXPHOS-related genes in the subtype 2 (Fig. 2A, B). Various stem cell populations are known for preferential utilization of glycolysis over mitochondrial oxidative metabolism as it allows them to be independent of oxygen level, and also to preserve the genomic integrity by reducing ROS. [65]. Aerobics metabolism and ROS regulation play a significant role in the stem cell fate change and differentiation [66] and is known to be the primary mechanism of ATP production in differentiated (somatic) cells [67].

### Subtype 2 of seminoma demonstrate molecular features specific for non-seminoma germ cell tumors

Seminoma cells (TCam-2 cell line) can be reprogrammed into an EC-like cell fate and further differentiate into mixed non-seminoma TGCT containing different TGCT histological components (EC, YST, seminoma, teratoma) [68]. To test whether the revealed subtype 2 of seminoma is in a transition stage towards more differentiated TGCTs, we analyzed the expression of signature genes known for other TGCT histological types [51, 69]. For this analysis we did not use any cutoff for the Log2FoldChange value, we picked signature genes which expression level was significantly different between subtypes base on adjusted p-value (P-value < 0.05) (Supplementary Table S4). We found that subtype 2 tumor samples had elevated level of non-seminoma signature genes when subtype 1 demonstrated seminoma features (Supplementary Table S4). The role of the defined genes in GCT cells is not clear, however some of them are important for tumorigenesis in other types of cancer [70–73].

A seminoma signature gene LZTS1 (Leucine Zipper Tumor Suppressor 1), has high level of expression in identified seminomas of subtype 1. This gene plays role in cell cycle control moderating the transition from late S to G(2)/M stage [74]. It was also shown that expression of LZTS1 correlates with SOX17 and both are important regulators of human pluripotent stem cell (hPSC) endoderm specification. Overexpression of LZTS1 in hPSCs leads to increased expression of SOX17 [75]. Though there are no information on LZTS1 role in seminomas, or other TGCTs, we know that activation of SOX17 in human embryonic stem cells determines their specification into primordial germ cells and is used as a key marker of the earliest PGCs [64]. Therefore, we suggest that LZTS1 may play a role in maintenance of early PGC pluripotency state in subtype 1 seminomas through a positive feedback loop with SOX17, and that can be related to an early PGC cell ancestry.

TGCT signature genes overexpressed in the subtype 2 of seminoma were characteristic for 3 major histotypes of non-seminomatous TGCTs, embryonal carcinoma (EC), yolk sac tumor (YST) and teratoma (Ter) (Supplementary Table S4). GAL and GPC4 gene signatures specific to EC and play an important role in embryonic development and pluripotency. Surface protein glypican 4 (Gpc4) is a component of the signaling machinery regulating embryonic stem cell (ESC) maintenance. In ESC, Gpc4 modulates the response to Wnt ligands and regulates activation of b- catenin signaling. GPC4 is important for teratoma lineage specification as loss of GPC4 in ESC makes ESC incompetent from developing teratomas [76]. Elevated expression of GPC4 may be a sign of EC-like stemness of seminoma subtype 2 with an intrinsic tendency to transform into multiple cell lineages including teratoma. Moreover, our data show that subtype 2 seminomas overexpress signature genes of more differentiated TGCTs, as YSTs (APOA2, BMP2, FAM89A, FOXA2, RAGE, VTN) and teratomas (MFAP4, NFKBIZ, and TSPAN8) supporting the hypothesis stated above. Importantly, we found that subtype 2 seminomas have increased expression of FOXA2 (Forkhead Box A2), a transcriptional factor that induces differentiation and microenvironment-triggered reprogramming of seminoma cells (TCam-2) into embryonal carcinoma [68]. FOXA2 is only upregulated for a limited amount of time and associated with a period of seminoma differentiation into non-seminoma lineages (EC). Once adaptation to the newly acquired cell fate is completed, FOXA2 expression is downregulated and it is not detectable in any TGCT subtypes [68]. The fact that we see elevated level of FOXA2 in subtype 2 seminoma supports the idea that this subtype is at an early transitional stage into EC fate. Therefore, histological examination has not revealed significant morphology differences between both seminoma subtypes. However, transcriptomic analysis detects significant changes in multiple molecular process including DNA repair and stemness maintenance. Several recent studies demonstrated that seminomas harbor certain molecular patterns which bring them closer to non-seminomas [77–80]. Moreover, a process of reprogramming of pure seminoma cells might take place, which results in the progression from seminoma to totipotent embryonal carcinoma cells that have the capacity of originating tumor components of all types of NSE GCTs [77, 80]. This process should be considered during the design of clinical therapeutic strategies and it increases the risk of poor prognosis [81, 82]. We hypothesize that revealed subtype 2 can be a precursor of mixed TGCT seminomas.

### Genomic and epigenomic features of seminoma subtype 2 revealed similarity with non- seminomatous GCTs

To further understand the biology of identified two subtypes of seminomas, we compared their genomic and epigenomic features. TGCTs have very low mutation burden, and only three genes were shown to contain recurrent somatic mutations: KIT, KRAS and NRAS [2]. We did not observe any association of a particular mutation pattern of these genes with the revealed seminoma subtypes. Nearly all TGCTs contain significant arm level gain of chromosome 12p (conventional marker of TGCT type II) and 21q, and moderate arm level gain of chromosomes 7 and 8 [2, 83]. Both identified seminoma subtypes revealed arm-level gain of all chromosomes mentioned above, as well as arm level loss of chromosomes 4, 5 and 11, with higher loss rate in subtype 1 (Supplementary Fig. S2).

TGCTs have a unique feature that distinguish them from other cancer types, this is highly recurrent chromosome arm level amplifications and loss of heterozygosity (LOH). It was found that TGCTs possess significant enrichment in number of chromosome arms with more than one allele amplified compared to 20 other cancer types [84]. Comparison of pure seminoma and seminoma originated from mixed TGCT demonstrated significantly higher LOH rate for the mixed seminoma [78]. We compared LOH data taken from TCGA PanCanAtlas study [53] for the identified seminoma subtypes. Our analysis revealed that for all chromosome arms except 9p and 11q LOH rate is higher for seminoma subtype 2 (Fig. 4A). Significant difference was detected for arms 3p, 4p, 6p, 7p, 11q, 12q, 15q, 17p and 20p. For the loci 13p, 14p, 15p, 21p and 22p no LOH was observed. Increased LOH rate of seminoma subtype 2 may be associated with impaired HR repair system, as we have noticed that in comparison to subtype 1, subtype 2 has decreased expression of genes associated with HR repair due to more advanced differentiation status. There are multiple studies on ovarian cancer that demonstrate that one of the reasons for increased LOH rate is deficiency of homologous recombination related to BRCA1 and BRCA2 mutation status [85–87].

**Figure 4.**
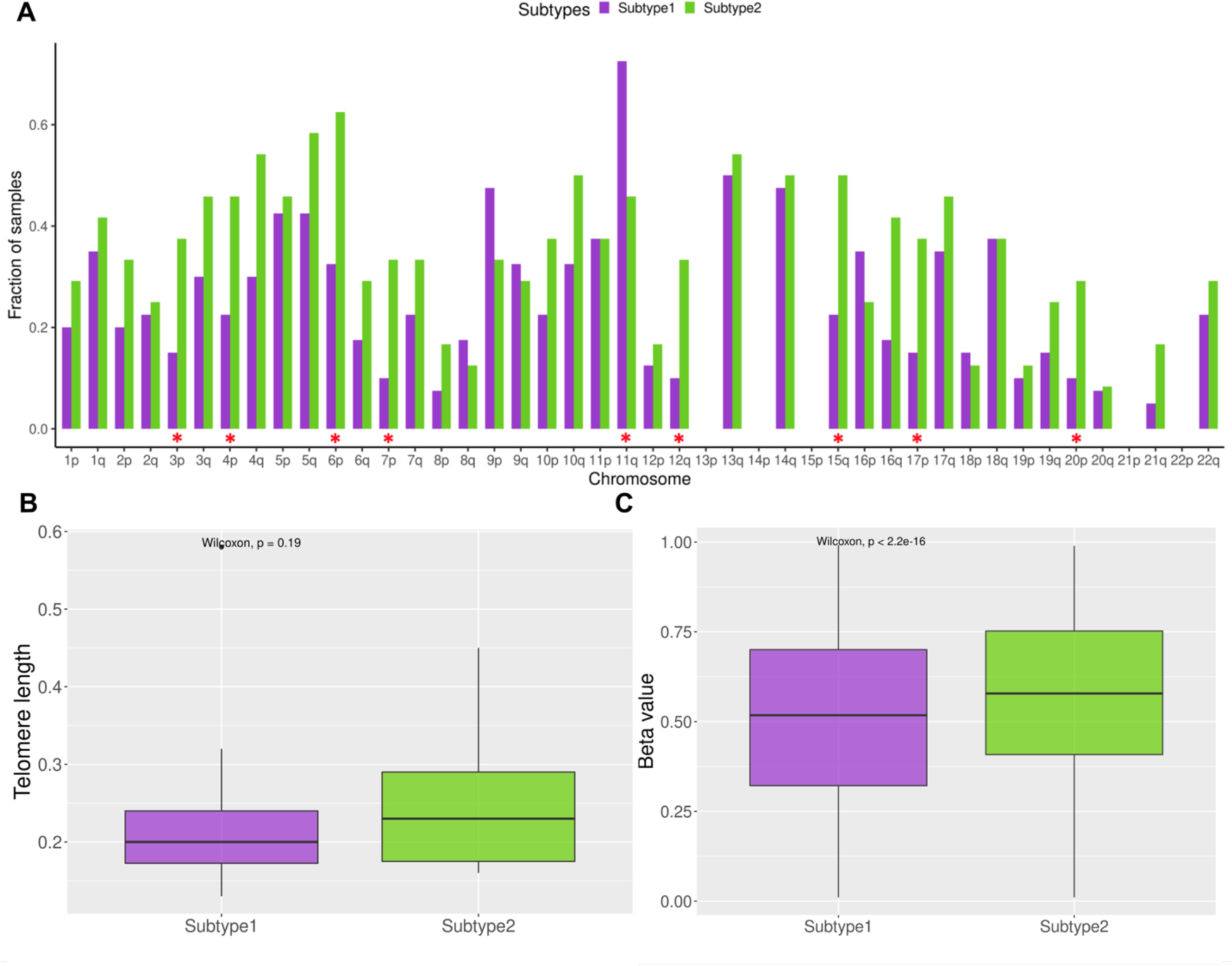
**(A)** Fraction of samples with observed LOH. Red asterisks denote significant difference between seminoma subtypes (Chi-squared test P-value < 0.05). **(B)** Boxplot of telomere length comparison between seminoma subtypes. **(C)** Boxplot of methylation level (beta value) comparison between seminoma subtypes.

Another molecular characteristic that differentiates SE from NSE is dominant telomere elongation in non-seminoma samples [88]. Our analysis revealed that telomere elongation is higher for subtype 2, however the difference is not significant (Fig. 4B). Moreover, among genes that have positive correlation between telomere elongation and gene expression in non-seminomas [88], we identified three genes which are significantly overexpressed in subtype 2: MT2A (Log2FC = 1.15, P-value = 3.9E-06), SLC16A3 (Log2FC = 1.1, P-value = 2.9E-04) and PDHA1 (Log2FC = 0.7, P-value = 6.2E-05).

Another genomic trait of non-seminomas versus seminomas is high level of DNA methylation [89–90]. Our analysis showed that subtype 2 of seminoma had significantly higher level of DNA methylation (Wilcoxon P-value < 2.2E-16) than subtype 1 (Fig. 4C). If we trace methylation status of TGCT precursor cell (PGC), we will find that at the earliest stage when ESC transforms into gonocyte (early PGC), it loses its DNA methylation pattern. In transition from early to late PGC and during further specialization of PGC into spermatogonia, cells will start *de novo* DNA methylation to re-establish the parental imprinting pattern [63, 91]. Thus, DNA methylation level of TGCT cells has strong association with their stemness and can reflect transition of TGCT tumor cells into more differentiated tumor subtype. It was demonstrated that during reprograming of seminoma cells (TCam-2) into EC, their global DNA methylation level strongly increases to a level of EC cells [92]. Therefore, we hypothesize that increased DNA methylation detected in subtype 2 seminomas is in a great alignment with previously discussed data, and it demonstrates more differentiated status of subtype 2 seminoma cells. It was shown that seminoma cell (TCam-2) methylation status has inverse correlation with resistance to cisplatin [90]. TCam-2 demethylation resulted in decreased resistance to cisplatin and increased expression of pluripotency markers (NANOG, POU5F1) and late PGC marker (VASA) [90].

### Long non-coding RNA expression pattern of seminoma subtype 2 suggests increased resistance to chemotherapy

Long non-coding RNAs (lncRNAs) were defined relatively recently as non-coding RNAs longer than 200 nucleotides. These molecules have been found to have crucial role in cancer utilizing large variety of functions including oncogenesis and tumor suppression and their function list is rapidly emerging [93]. TGCTs are not the exception to this rule, expression levels of several oncogenic lncRNAs have been associated with germ cell tumors [94]. Unsupervised clustering of lncRNAs genes based on their expression level showed two large clusters which are very similar to identified subtypes based on the whole transcriptome (Fig. 5A). LncRNAs play crucial role in the development of cisplatin resistance [95]. TGCTs are highly sensitive to chemotherapy and radiotherapy. Interestingly that seminomas are more sensitive than non-seminomas, that might be related to their differentiation status (mature teratomas are the most chemotherapy resistant TGCTs) and active DNA repair mechanisms [96, 97]. Similar pattern of drug resistance increment was noticed during the development of sperm from PGCs, and therefore it was proposed that seminomas and non-seminomas are derived from cells at different stages of germ-cell differentiation [96]. We identified that five lncRNAs responsible for cisplatin resistance in different cancer types are overexpressed in seminoma subtype 2 (Fig. 5B). H19 was shown to utilize pro-tumorigenic function and promote cisplatin resistance in TGCTs [95, 98]. Other four lncRNAs (NEAT1, PVT1, SFTA1P, TRPM2-AS) are responsible for cisplatin resistance in lung, gastric and ovarian cancers [95]. Advanced differentiation status, HR repair deficiency, increased methylation status and overexpression of lncRNA associated with cisplatin resistance allow us to hypothesize that revealed subtype 2 of seminoma is more resistant to genotoxic drugs. Therefore, patients with subtype 2 seminoma may require adjustments of a treatment protocol or development of alternative treatment approaches.

**Figure 5.**
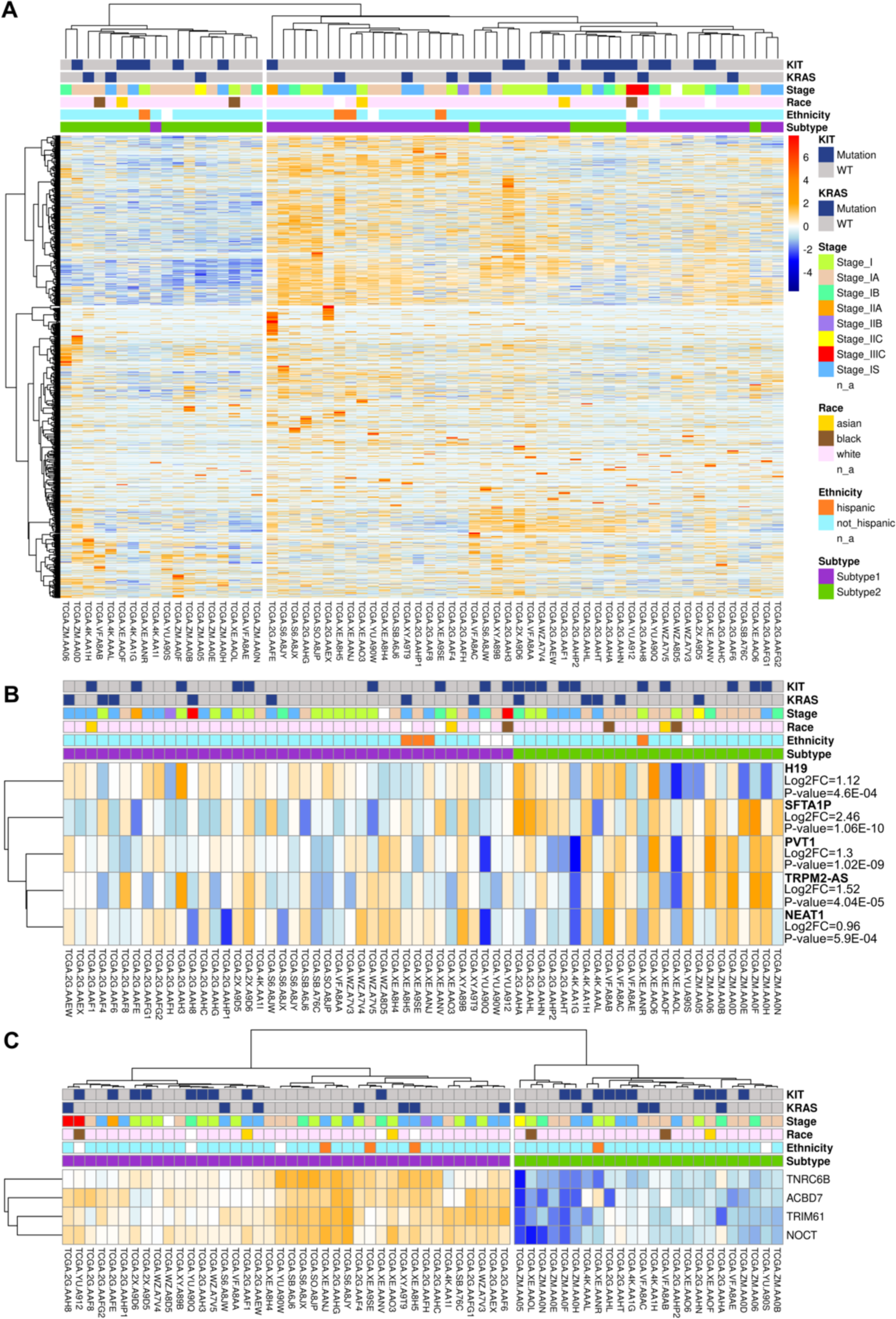
**(A)** Unsupervised hierarchical clustering of seminoma samples based on transcriptomic data of lncRNAs. **(B)** Five lncRNAs that promote cisplatin resistance and are overexpressed in subtype 2. **(C)** Potential biomarkers for seminoma subtypes identification (overexpressed in subtype 1).

### Potential biomarkers for histological differentiation of seminoma subtypes

We identified four potential biomarkers which are capable to distinguish two seminoma subtypes (Fig. 5C; Supplementary Table S5). All these genes are overexpressed in subtype 1 and show specificity no less than 79%, which is in the specificity range of clinically used biomarkers for various cancer types [99]. Overexpressed genes in subtype 2 do not show high enough specificity, so we do not consider them. All of the identified biomarkers were assessed in other cancer types, but were not previously evaluated for TGCTs. Nocturnin (NOCT) shows the highest specificity level of 92%. NOCT is overexpressed in squamous cell lung cancer and has potential as biomarker for this type of cancer [100]. TNRC6B shows specificity of 87.5%. Alterations in expression level of this gene was shown to contribute to carcinogenesis [101]. Finally, TRIM61 and ACBD7 show specificity of 79.2%. TRIM61 was previously suggested as prognostic biomarker for lung squamous cell cancer [102], while ACBD7 is overexpressed in Hürthle cell carcinoma [103]. Combination of potential biomarkers NOCT and TNRC6B results in the highest specificity of 96%. Discussed potential biomarkers were defined using computational analysis and require further experimental validation.

Summing up, our analysis showed that pure seminoma cases can be further subdivided into 2 main subtypes. The subtypes differ in 1) pluripotency stage, 2) activity of DSB DNA repair mechanisms, 3) rate of LOH, DNA methylation and telomere elongation, 4) expression of lncRNA associated with cisplatin resistance. In comparison to more pluripotent seminoma subtype 1, seminoma subtype 2 shows signs of differentiation into non-seminoma TGCT and may have higher resistance to platinum-based chemotherapy.

## Discussion

TGCTs represent highly heterogeneous group of cancers, starting with broad separation of seminoma and NSE GCTs [2]. Moreover, each TGCT histological type has demonstrated existing internal heterogeneity leading to diverse patient outcomes [96]. Our analysis of the dominant TGCT subtype seminoma, demonstrated that on the transcriptomic level seminoma samples can be further classified into 2 subtypes. Subtype 2 revealed several molecular features that can be associated with undergoing differentiation of subtype 2 into non-seminomatous lineages through the stage of embryonal carcinoma (Fig. 6). It is important to consider that seminoma subtype 2 showed impaired HR repair in comparison to subtype 1. Subtype 2 depends on more efficient c-NHEJ repair mechanism that allows these tumors to repair DSBs in a timely manner avoiding apoptotic cell death, which may potentially explain greater resistance to chemotherapy. Analysis of patient outcome has not revealed significant difference between seminoma subtypes after initial treatment. However, detailed evaluation considering amount and duration of platinum-based therapy and long-term outcomes are necessary for the final conclusion. Hypothetically, subtype 2 seminoma cells may be responsible for seminoma recurrence after chemotherapy through mechanisms of undertreatment or cisplatin-resistance. Relapsed patients have a 50% chance of disease-specifically mortality; and salvage treatment drastically worsen side effect profiles. However, some patients suffered progressive cancer disease despite high-dose chemotherapy [104]. In addition, platinum-base chemotherapy, which is used for TGCTs treatment, significantly decreases patients’ quality of life and can cause complex of around 40 severe long-term side effects including secondary cancers and death [17]. Circulating platinum concentration can remain up to 1,000 times above the normal level for 20 years after the chemotherapy completion and is associated with many delayed side-effects [105]. Therefore, it is important to develop fewer toxic solutions for TGCT therapy. Molecular differences identified between 2 seminoma subtypes will potentially lead to important practical applications. There are multiple studies showing that PARP inhibitors (FDA approved for various ovarian and breast cancers) can be efficiently used against HR-deficient tumors as it induces DSB formation and in addition inhibits alternative repair mechanisms such as microhomology-mediated end joining (MMEJ) [106, 107]. Our analysis showed that subtype 2 seminoma has deficiency in HR repair that makes it a suitable candidate for PARP inhibitor therapy or combination therapy of PARP inhibitors with platinum compounds. However, in some tumors PARP inhibitors may elicit a tumoricidal effect by enhancing c-NHEJ [106, 107], therefore preliminary in vitro studies are required. On the contrary to subtype 2, subtype 1 has increased activity of HR repair that also can be used for a new therapy development. For example, tyrosine kinase inhibitor erlotinib disrupts nuclear function of BRCA1 and attenuates HR activity. As the result, it causes sensitization of cancer cells to radiation therapy [108, 109]. There is also evidence that proteasome inhibitors targeting HR proteins can cause cisplatin sensitization [110].

**Figure 6.**
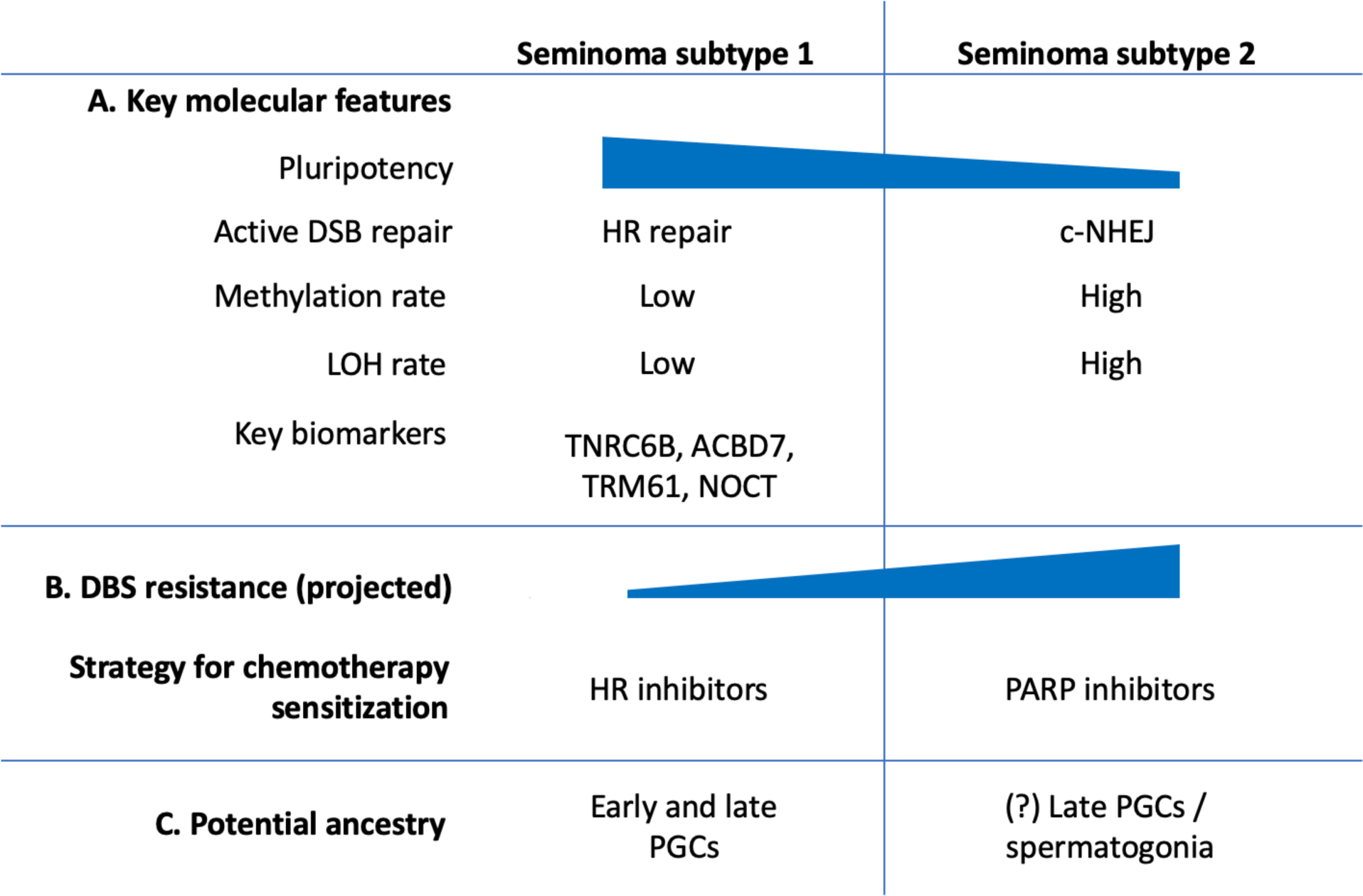
Main features of seminoma subtypes 1 and 2.

## Conclusions

Computational analysis of omics data of pure seminoma samples from TCGA revealed two distinct seminoma subtypes. Subtype 1 has higher pluripotency rate and demonstrated signs of elevated HR repair activity. On the contrary, subtype 2 showed features of more differentiated cell type and resembles non-seminoma TGCTs (overexpression of signature genes, increased DNA methylation, high rate of LOH and elongated telomers). We also detected that subtype 2 samples had increased expression level of lncRNAs responsible for cisplatin resistance in different cancer types including TGCTs. We hypothesize that drugs targeting HR repair (subtype 1) and other DSB repair mechanisms as c-NHEJ and MMEJ (subtype 2) can significantly increase sensitivity of revealed seminoma subtypes to chemotherapy and irradiation, though in vitro and in vivo studies are required to support the hypothesis. Development of seminoma subtype-specific therapy can help to overcome chemotherapy overtreatment in TGCT patients and improve quality of life for TGCT survivors.

## CRediT Author Contribution

**Kirill E. Medvedev:** Conceptualization, Methodology, Software, Validation, Formal analysis, Investigation, Data Curation, Visualization, Writing - Original Draft, Project administration. **Anna V. Savelyeva**: Conceptualization, Validation, Formal analysis, Investigation, Writing - Original Draft. **Aditya Bagrodia:** Resources, Funding acquisition, Writing - Review & Editing. **Liwei Jia:** Investigation, Writing - Review & Editing. **Nick V. Grishin:** Conceptualization, Resources, Funding acquisition, Writing - Review & Editing.

## Acknowledgements

Authors are grateful to TCGA data portal for providing access to TGCT datasets.

