## Supplementary Figures S1-S2, Tables S4-S5 for "Integrated molecular analysis reveals two distinct subtypes of pure seminoma of the testis"

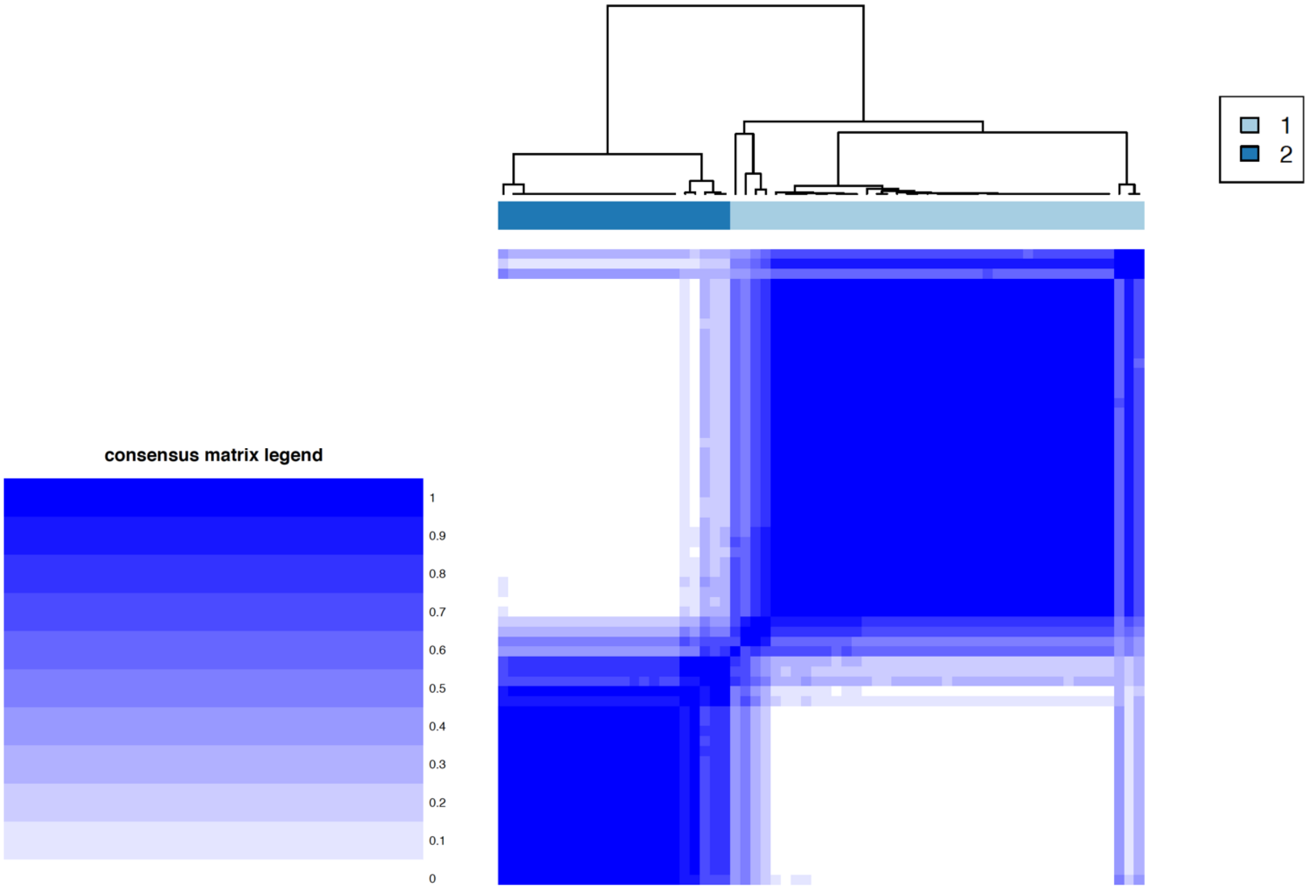


**Fig. S1. Consensus matrix plot of 64 seminoma samples based on transcriptomic data.**

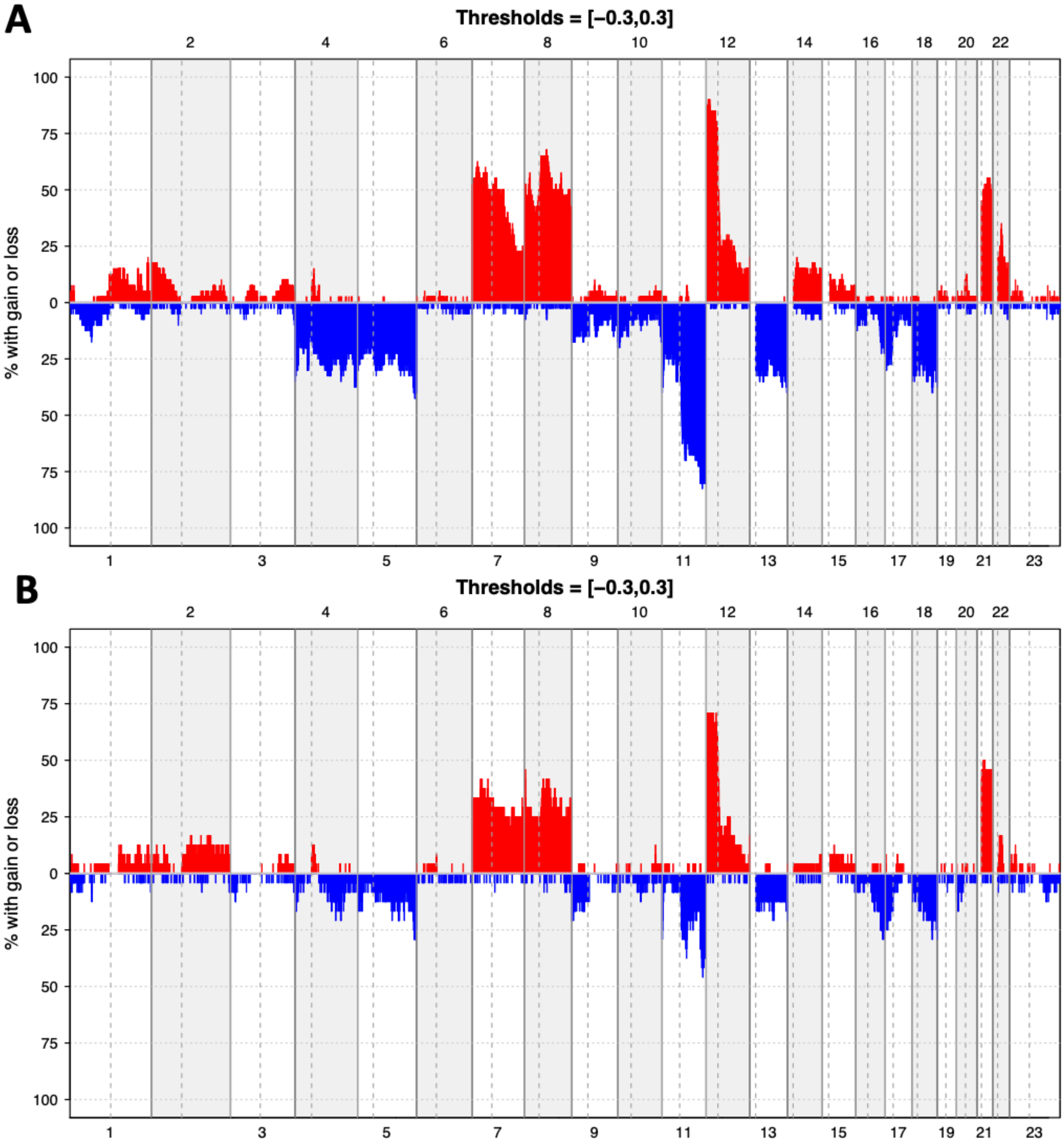


**Fig. S2. Chromosome copy number gains (red) and losses (blue) for seminoma (A) subtype1 and (B) subtype2.**

| Signature genes | Log2FC | Adjusted P-value | Seminoma subtype overexpression |
|---|---|---|---|
| Seminoma | | | |
| LZTS1 | 0.71 | 0.00063 | Subtype1 |
| Embryonal carcinoma | | | |
| GAL | 1.16 | 0.049 | Subtype2 |
| GPC4 | 1.2 | 0.0025 | Subtype2 |
| Yolk sac tumor | | | |
| APOA2 | 1.31 | 0.01 | Subtype2 |
| BMP2 | 0.85 | 0.0059 | Subtype2 |
| FAM89A | 0.88 | 0.0002 | Subtype2 |
| FOXA2 | 1.02 | 0.036 | Subtype2 |
| RAGE | 1.25 | 1.97E-09 | Subtype2 |
| VTN | 1.53 | 0.0002 | Subtype2 |
| Teratoma | | | |
| MFAP4 | 1.01 | 0.069 | Subtype2 |
| NFKBIZ | 0.78 | 0.0007 | Subtype2 |
| TSPAN8 | 2.36 | 2.06E-08 | Subtype2 |

**Table S4. Signature genes of different TGCT histological types overexpressed in seminoma subtypes.**

| Gene | baseMean | Log2FC | Adjusted P-value | Specificity as biomarker |
|---|---|---|---|---|
| NOCT | 618.28 | 1.05 | 5.74E-12 | 92% |
| TRIM61 | 121.91 | 1.19 | 4.37E-11 | 79.2% |
| ACBD7 | 63.14 | 2.05 | 1.12E-09 | 79.2% |
| TNRC6B | 3319.48 | 1.03 | 5.07E-13 | 87.5% |

**Table S5. Potential biomarkers for seminoma subtype 1 identification.**
